# Regulation of mannitol metabolism in *Enterococcus faecalis* and association with *par*_EF0409_ toxin-antitoxin locus function

**DOI:** 10.1101/2022.02.04.479212

**Authors:** Srivishnupriya Anbalagan, Jessie Sadlon, Keith Weaver

**Affiliations:** Division of Basic Biomedical Sciences. Sanford School of Medicine. University of South Dakota, Vermillion, SD, USA

**Keywords:** *Enterococcus faecalis*, mannitol metabolism, phosphotransferase systems, toxin-antitoxin-systems

## Abstract

The *par*_EF0409_ type I toxin-antitoxin locus is situated between genes for two paralogous mannitol-family phosphoenolpyruvate phosphotransferase systems (PTS). In order to address the possibility that *par*_EF0409_ function was associated with sugar metabolism, genetic and phenotypic analyses were performed on the flanking genes. It was found that the genes were transcribed as two operons; the downstream operon essential for mannitol transport and metabolism and the upstream operon performing a regulatory function. In addition to genes for the PTS components, the upstream operon encodes a gene similar to *mtlR*, the key regulator of mannitol metabolism in other Gram-positive bacteria. We confirmed that this gene is essential for regulation of the downstream operon and identified putative phosphorylation sites required for carbon catabolite repression and mannitol-specific regulation. Genomic comparisons revealed that this dual operon organization of mannitol utilization genes is uncommon in enterococci and that association with a toxin-antitoxin system is unique to *E. faecalis*. Finally, we consider possible links between *par*_EF0409_ function and mannitol utilization.

**Importance:** *Enterococcus faecalis* is both a common member of the human gut microbiota and an opportunistic pathogen. Its evolutionary success is partially due to its metabolic flexibility: in particular, its ability to import and metabolize a wide variety of sugars. While a large number of phosphoenolpyruvate phosphotransferase sugar transport systems have been identified in the *E. faecalis* genome bioinformatically, the specificity and regulation of most of these systems remains undetermined. Here we characterize a complex system of two operons flanking a type I toxin-antitoxin system required for the transport and metabolism of the common dietary sugar mannitol. We also determine the phylogenetic distribution of mannitol utilization genes in the enterococcal genus and discuss the significance of association with toxin-antitoxin systems.

## Introduction

Enterococci are notorious for their metabolic versatility and intrinsic resistance to inhospitable environments. One aspect of that versatility is the ability to transport and utilize a variety of carbohydrates as carbon sources. At least 13 sugars are metabolized by all enterococci and over 30 are used by at least two species (1). Import of most of these sugars is accomplished by phosphoenolpyruvate phosphotransferase systems (PTS) that couple uptake with phosphorylation of the substrate. The *Enterococcus faecalis* strain OG1RF encodes 39 predicted PTS systems based on NCBI genome annotation, but the targets for most of these transporters have yet to be determined experimentally. In addition, genome sequencing indicates that a number of PTS systems are encoded on mobile genetic elements (MGE) leading to extensive within species variability and a possible association with virulence (2). Indeed, PTS systems have been associated with increased onset of colitis in a mouse model (3), increased killing of macrophages (3), enhanced stress resistance (4), endocarditis and biofilm formation (5), and murine intestinal colonization (6) in *E. faecalis* and *E. faecium*.

Mannitol is an abundant sugar found naturally in plants and is used in food as an artificial sweetener particularly in candies, dried fruits and chewing gum and is poorly absorbed by the intestines (7). Thus, enterococci are very likely to find it as a common carbon source in the intestinal tract where it resides. An *E. faecalis* encoded mannitol-specific PTS component, designated *mtlF*, and an associated mannitol-1-phosphate dehydrogenase gene, *mtlD*, were originally cloned and characterized by Fischer *et al* in 1991 (8). However, sequencing of the OG1RF genome revealed a complex organization of two paralogous PTS systems: an upstream cluster that also included a putative mannitol-responsive positive transcriptional regulator, *mtlR*, and a downstream cluster that included *mtlD* (Fig. 1). It was the latter cluster that had been previously cloned by Fischer *et al*. Between the two clusters is a type I toxin-antitoxin (TA-1) system, designated *par*_EF0409_ (9), related to *par*_pAD1_ originally described on the *E. faecalis* pheromone-inducible conjugative plasmid pAD1 (10). Toxin-antitoxin (TA) systems are ubiquitous on bacterial MGE where they function to facilitate stable inheritance by a post-segregational killing (PSK) mechanism (11, 12). They are classified based on the nature and mechanism of action of their antitoxins; the antitoxins of TA-1 systems are regulatory small RNAs (sRNA) that bind to and prevent translation of the toxin mRNA (13-15). The antitoxin sRNA is less stable than the toxin mRNA so plasmid loss leads to selective degradation of the sRNA and translation of the toxin, thereby inhibiting growth of plasmid-free segregants. The pAD1-encoded *par*_pAD1_ was shown to function in just such a manner (16-18). The explosion of genomic sequencing in bacteria revealed that paralogs of many plasmid-encoded TA systems were also present on bacterial chromosomes (19). The function of these chromosomally-encoded systems are controversial and mostly unknown (15, 20, 21). The function of *par*_EF0409_ is similarly obscure.

**Figure 1.**
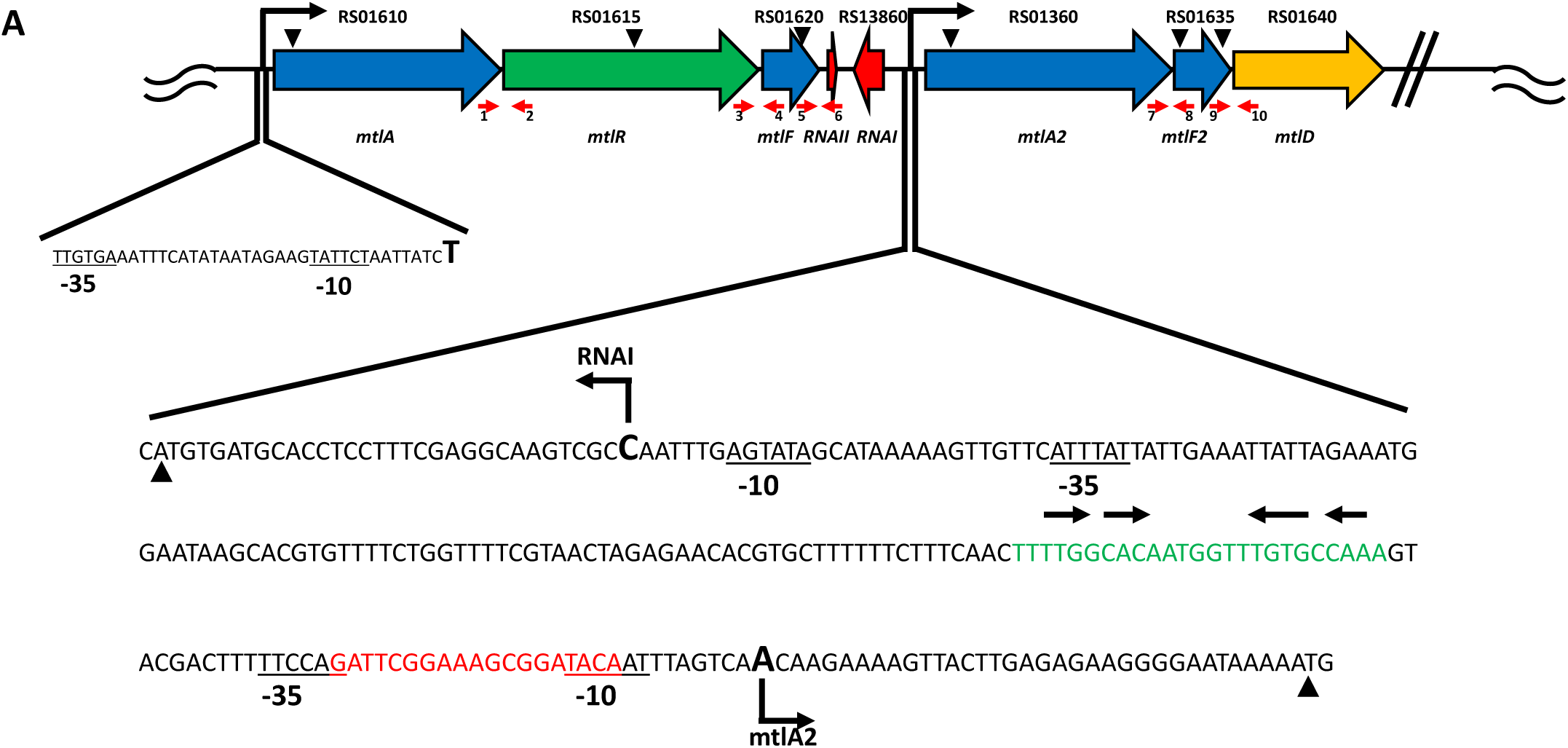
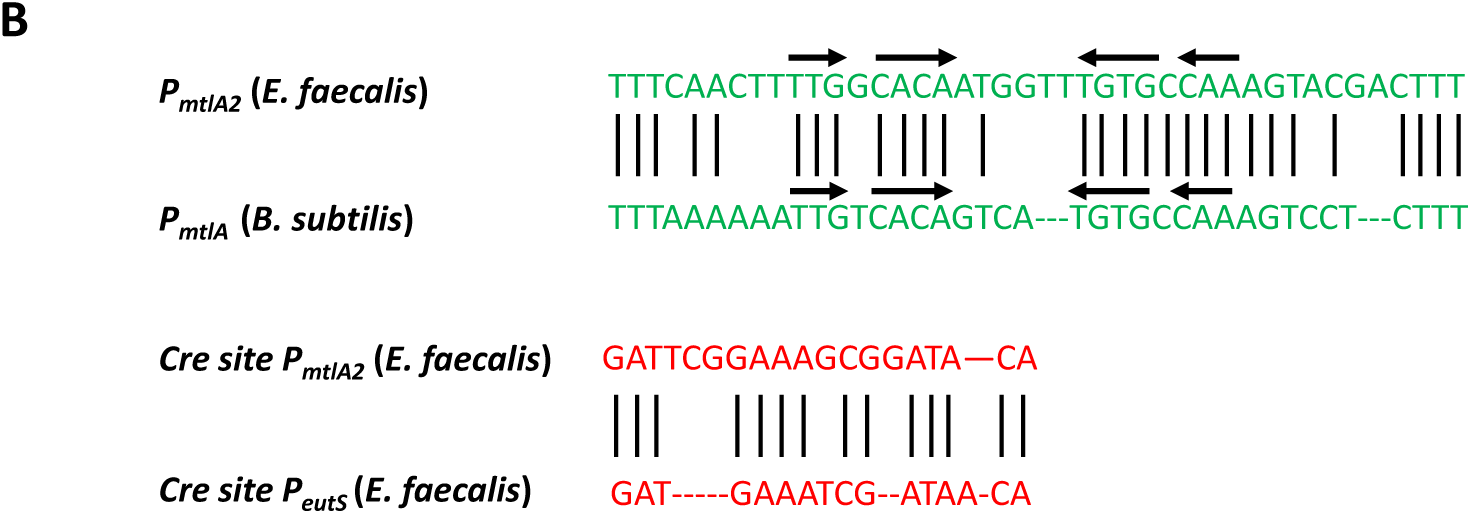
**A. Genetic and transcriptional organization of the *mtlA operon, par*_*EF0409*_ locus, and *mtlA2* operon.** Large arrows depict the position and direction of transcription of the relevant genes color coded for function. Blue: PTS transport. Orange: metabolism. Green: regulation. Red: *par*_EF0409_ TA-1. Current NCBI OG1RF numerical gene designations are shown above the genes while the orthologous gene designation based on sequence homology is shown below. Small inverted triangles above each gene indicate transposon insertion sites. Broken arrows represent promoters for the *mtlA* and *mtlA2* operons with expanded sequence showing transcription start sites identified by 5’RACE. Predicted MtlR and CcpA binding sites are in green and red, respectively. Translational start sites of *mtlA2* and *fst* are marked with a filled triangle. Small red arrows at the base of each gene represent the primer sites used for RT-PCR to determine operon organization. Numbers under the small arrows correspond to the primer sequence in Table*. **B. Alignment of predicted MtlR binding and *cre* sites**. MtlR binding site is compared to previously determined *B. subtilis* MtlR binding site (28) and *cre* site as determined for the *E. faecalis eutS* gene (29).

In this report, we characterize the complex organization of the mannitol utilization genes and determine their functional roles. We demonstrate that 1) the genes are arranged in two operons that flank *par*_EF0409_, 2) the PTS system encoded in the downstream operon is the primary transport system for mannitol metabolism, 3) the putative transcriptional regulator, MtlR, positively regulates the downstream genes and phosphorylation sites shown to be critical for regulation in orthologs are important for its function, and 4) the PTS system encoded by the upstream operon plays an auxiliary regulatory role, presumably by modulating MtlR function. We also present several lines of circumstantial evidence that the function of *par*_EF0409_ relates in some yet undefined way to regulation of mannitol metabolism.

## Results

### Functional analysis and transcriptional organization of the *mtlA* and *mtlA2* operons

In order to evaluate the functional roles of the putative *mtlA* and *mtlA2* operons, transposon insertion mutants in *mltA, mtlR, mtlF, mtlA2*, and *mtlF2* (see Fig. 1) were obtained from the ordered transposon mutant library produced by the laboratory of Dr. Gary Dunny (22, 23). All transposon insertion strains grew normally on glucose (M9YEG) but showed varying levels of defects in growth on mannitol (M9YEM), supporting a role for these genes in mannitol metabolism. Insertions in *mtlR, mtlA2*, and *mtlF2* failed to grow on mannitol, while those in *mtlA* showed reduced growth (Fig. 2A), and insertions in *mtlF* had no effect (not shown).

**Figure 2.**
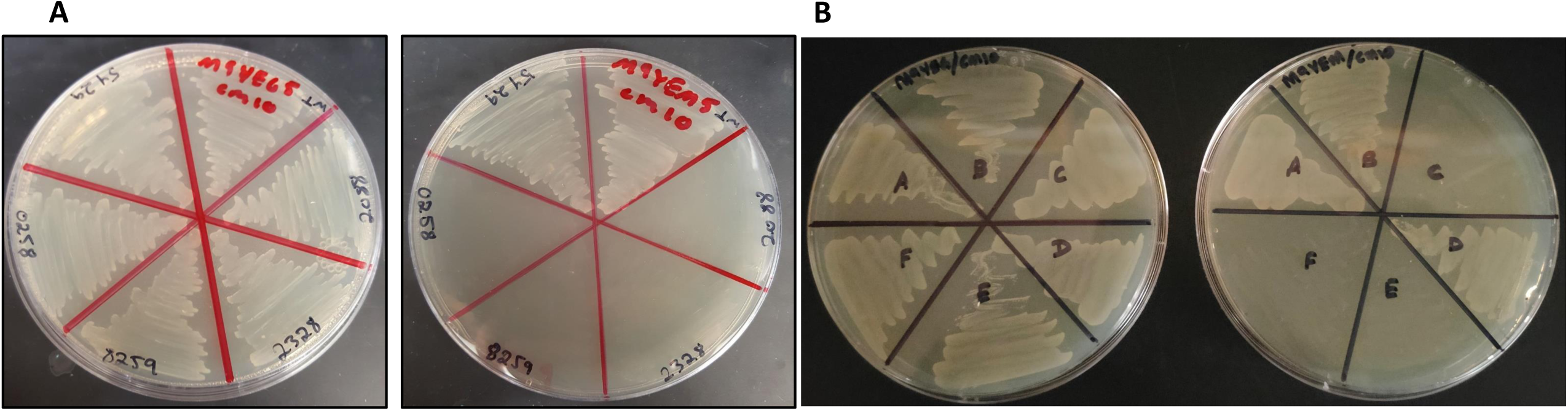
**A**. Growth of mutants with transposon insertions in the *mtlA* and *mtlA2* operons on M9YEG (left) and M9YEM (right): Clockwise from top right: OG1RF (WT), *mtlF2* (2088 & 2328), *mtlR* (8259), *mtlA2* (0258), and *mtlA* (5429). **B**. Complementation of OG1RF mutants with ectopic *mtlR*. Cells were grown on M9YEG (left) and M9YEM (right). A: OG1RF (pCIE). B: OG1RF (pCIE::*mtlR*). C: OG1RF*ΔmtlR* (pCIE). D OG1RF*ΔmtlR* (pCIE*::mtlR)*. E: OG1RF*Δ*MBS (pCIE). F: OG1RF*Δ*MBS (pCIE*::mtlR)*.

The genetic organization of the relevant genes suggested transcription as two operons: *mtlA-mtlR-mtlF* (*mtlA* operon) and *mtlA2-mtlF2-mtlD* (*mtlA2* operon) (Fig. 1). Promoters for the putative operons were identified by 5’-RACE using primers within *mtlA* and *mtlA2* (Table S1) on RNA purified from cells grown in either glucose or mannitol. Each primer produced a single product (Fig. 3) and the transcriptional start sites identified along with putative promoter sequences are shown in Fig. 1A. Identical transcription start sites were mapped in glucose and mannitol-grown cells, but substantially more product was obtained for *mtlA2* in mannitol-grown cultures (Fig. 3). 5’-RACE performed with primers within *mtlR, mtlF, mtlF2*, and *mtlD* produced no detectable products (data not shown) suggesting that these genes were transcribed from upstream promoters. To confirm the operon structure, RT-PCR was performed on the same RNA samples with primers straddling the junction between genes of interest (Fig. 1 and Table S1). Products of appropriate size were found between all genes suspected of belonging to either the *mtlA* or *mtlA2* operons but not in controls lacking reverse transcriptase (Fig. S1 and S2). Interestingly, a product was also obtained with primers flanking the *mtlF* and RNA II_EF0409_ junction suggesting that, in addition to transcription from its own promoter, transcriptional readthrough from the upstream operon might affect RNA II_EF0409_ antitoxin levels. The pattern of RNA-seq reads was also consistent with an *mtlARF* and *mtlA2F2D* organization (data not shown). As shown in Fig. 1, the *mltA2* and RNA I_EF0409_ divergent promoters are separated by only 144 nt and most likely contain at least one transcription factor binding site (see below) suggesting possible coordination of expression.

**Figure 3.**
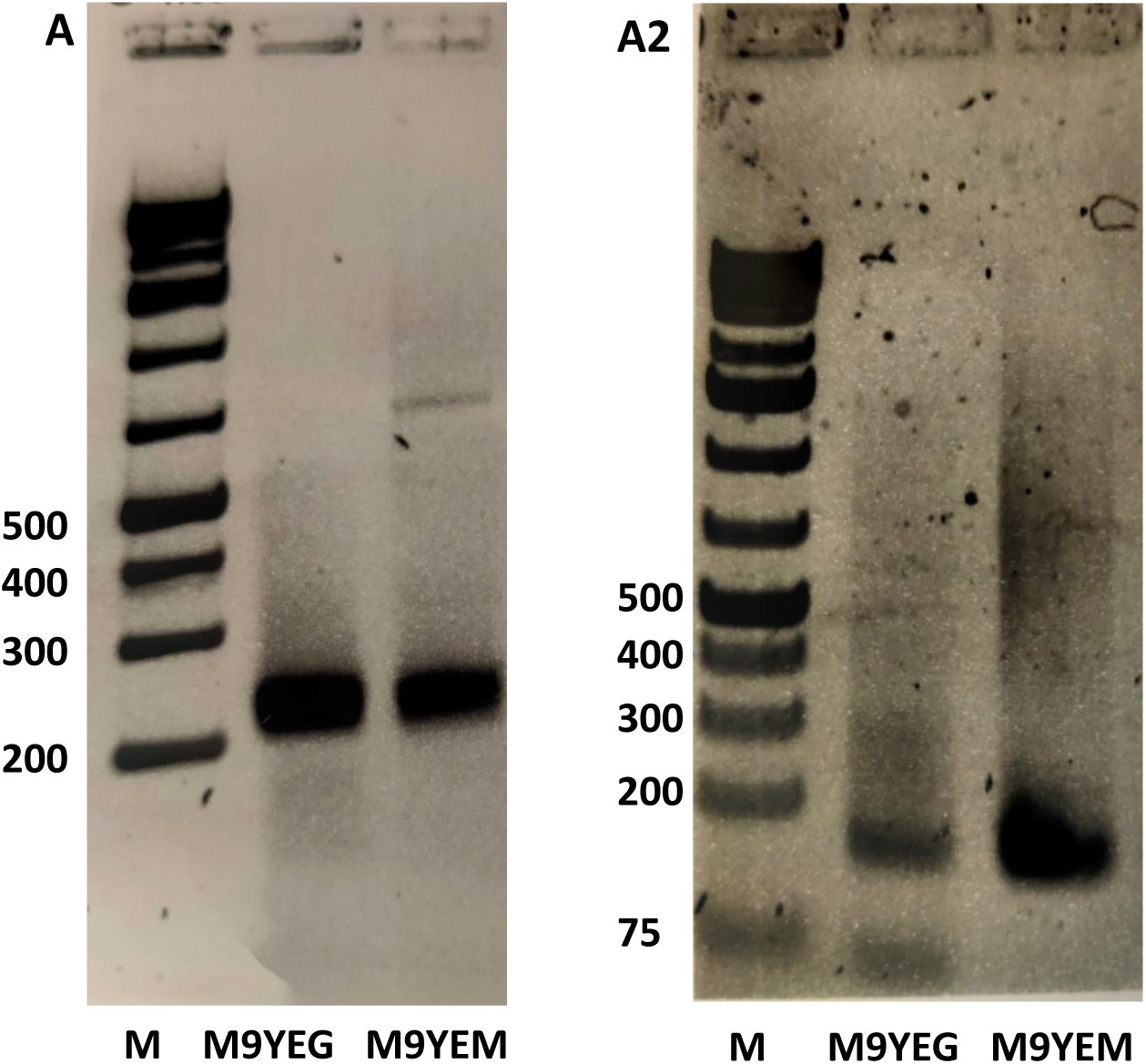
5’-RACE results with primers specific for *mtlA* **(A)** and *mtlA2* **(A2)** in glucose and mannitol. Molecular weight markers (M) are on the left of each gel. Transcriptional start sites determined by sequencing these products are shown in Fig. 1A.

Fischer *et al* (8) previously suggested that *mtlF* (*mtlF2* here) and *mtlD* were co-transcribed based on the overlap of termination and initiation codons and the presence of an intrinsic terminator downstream of *mtlD*. However, their sequence showed an approximately 60 nt gap between *mtlA2* and *mtlF2* leaving in question the relationship of these two genes. Comparison of their sequence with the NCBI OG1RF sequence revealed several discrepancies, most critically an insertion of a T at position 309 of the Fisher *et al* sequence (OG1RF 311902) that changes the reading frame and results in premature termination of *mtlA2*. In the NCBI sequence only 14 nt separates the 3’ end of *mtlA2* from the 5’ end *mtlF2* and there is no intrinsic terminator between, consistent with the operon organization proposed here.

To resolve questions concerning potential polar effects of the transposon insertions, in-frame deletions of *mtlA, mtlR, mtlF*, and *mtlA2* were constructed. Growth curves of these constructs were performed in M9YEG and M9YEM. Growth of *ΔmtlR* was significantly impaired in M9YEM as compared to WT (Fig. 4). Indeed, growth of *ΔmtlR* on M9YEM was indistinguishable from growth on M9YE without added sugar (Fig. S3) suggesting that the mutant was unable to use mannitol for growth. The residual growth observed was likely due to utilization of other nutrients in the yeast extract and casamino acids in M9YEM. By contrast, the *ΔmtlA2* mutant continued to grow slowly after growth of *ΔmtlR* had ceased (Fig. 4). Growth of *ΔmtlR* was also slightly reduced in M9YEG while *ΔmtlA2* had no effect on growth in this medium. The growth of *ΔmtlA* was indistinguishable from WT in both M9YEG and M9YEM (data not shown). These results suggested that *mtlA2-mtlF2* encoded the EIICBA PTS components essential for mannitol-specific transport and *mtlR* encoded a positive regulator of the *mtlA2* operon.

**Figure 4.**
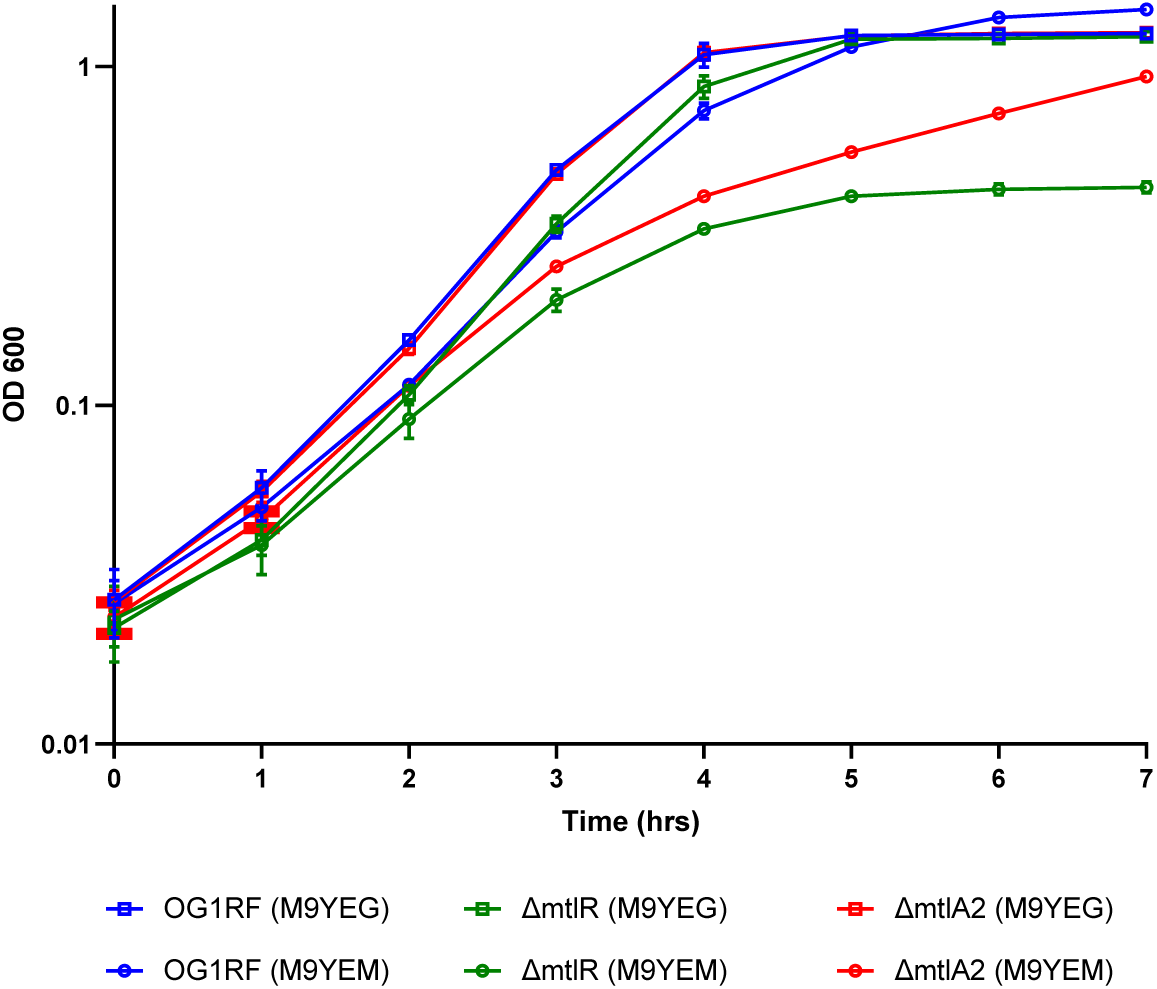
Growth of in-frame deletion mutants of *mtlR* and *mtlA2* on glucose (M9YEG) and mannitol (M9YEM). Results are an average of three independent experiments and standard deviation is shown in error bars. Error at later time-points was so small that the error bars are not visible beyond the marker.

### Regulation of the *mtlA* and *mtlA2* operons in response to mannitol

In order to investigate the transcriptional response to mannitol, RNA-seq was performed on WT cells grown in M9YEG and M9YEM. The three genes of the *mtlA2* operon were all induced greater than 100-fold when grown in M9YEM and were by far the most highly induced genes during growth on mannitol (Table 1 and supplemental). Levels of *mtlA* operon expression were not significantly different on glucose and mannitol. Results with *mtlA2* (Table 2) and *mtlA* (data not shown) were confirmed by qRT-PCR.

**Table 1.** Impact of growth in mannitol and deletion of *mtlR* in glucose on the expression of genes in the *mtlA* and *mtlA2* operons.

| Gene name | WT mtl vs. glu fold change | ΔmtlR vs. WT glu fold change |
| --- | --- | --- |
| *mtlA* | 0.9 | 0.7 |
| *mtlR* | 0.7 | NA<sup>b</sup> |
| *mtlF* | 1.4 | 1.4 |
| *mtlA2* | 311<sup>a</sup> | 0.2<sup>a</sup> |
| *mtlF2* | 257<sup>a</sup> | 0.2 |
| *mtlD* | 131<sup>a</sup> | 0.4 |

<sup>a</sup> Differences determined to be statistically significant
<sup>b</sup> NA = not applicable. Since the *mtlR* gene was deleted in this strain no reads were obtained.

**Table 2.** *mtlA2* levels in *mtlA* operon and MtlR binding site mutants

| Strain | Glucose | Mannitol |
| --- | --- | --- |
| WT | 0.228±0.046 | 2.603±0.047 |
| ΔmtlA | 0.771±0.015 | 2.549±0.059 |
| ΔmtlF | 0.495±0.026 | 2.527±0.033 |
| ΔmtlR | 0.059±0.017 | 0.078±0.013 |
| ΔMBS | 0.087±0.014 | 0.109±0.006 |

RNA-seq identified 217 other genes significantly altered in expression more than two-fold, 147 induced and 70 repressed (supplemental information). Among the most strongly affected genes were several that would be expected to impact the flow of carbon through the glycolytic cycle including glucosamine 6-phosphate deaminase, fructose bisphosphatase, and acetaldehyde CoA alcohol dehydrogenase (all induced over ten-fold), the putative glycerol dehydrogenase/dihydroxyacetone kinase operon (induced over six-fold) and beta-phosphoglucomutase (repressed approximately 22-fold). In addition, a number of ABC and sugar transport systems were moderately induced, perhaps because of the release of carbon catabolite repression. PTS systems orthologous to those for trehalose and mannose were among the most highly repressed genes. Since trehalose has been shown to induce beta-phosphoglucomutase in other organisms (24, 25) expression of these two genes may be linked.

The paralogous PTS system encoded in the *mtlA* operon was not essential for or induced by growth in mannitol, but the location of the *mtlR* gene within this operon suggested that it performed a regulatory function. The *mtlR* gene encodes a member of the BglG family of regulatory proteins and is most similar to the subfamily that acts as transcriptional activators (26). The growth defect of *ΔmtlR* in M9YEM suggested that the gene encoded a transcriptional activator of the *mtlA2* operon. Consistent with this hypothesis, qRT-PCR experiments showed that Δ*mtlR* negated mannitol induction of *mtlA2* (Table 2). Unexpectedly, Δ*mtlR* also reduced *mtlA2* expression in M9YEG; *mtlA2* expression was reduced four- and 33-fold in M9YEG and M9YEM, respectively, compared to WT control (Table 2). Because Δ*mtlR* also showed a growth defect in M9YEG, RNA-seq was performed to determine if transcription of any other genes were altered during growth in glucose. As expected, reductions were observed in all three genes of the *mtlA2* operon compared to WT (Table 1), although only the change in *mtlA2* levels was significant because of low basal levels of expression of *mtlF2* and *mtlD* on M9YEG. Only two other genes showed a significant change in expression over two-fold in Δ*mtlR* grown in M9YEG relative to WT: *dltX* involved in alanylation of teichoic acids and an orphan EIIC PTS component OG1RF_RS01210. Expression of both was increased approximately 2.5-fold in the Δ*mtlR* strain. It is possible that *mtlR* deletion resulted in post-transcriptional changes, e.g., protein phosphorylation, that affected growth in glucose.

Complementation of Δ*mtlR* with an ectopic copy of *mtlR* in the pCIE expression vector (9) confirmed the role of MtlR as an activator of *mtlA2* transcription. Complementation restored growth of Δ*mtlR* on M9YEM and WT *mtlA2* mRNA levels on both M9YEG and M9YEM (Fig. 2B and Table 3). Complementation was complete without cCF10 induction of the P_Q_ promoter, indicating that basal levels of transcription produced sufficient MtlR for a response to mannitol. P_Q_ induction produced only a small increase in *mtlA2* expression in M9YEM. In contrast, cCF10 induction of *mtlR* expression in M9YEG resulted in a six-fold higher transcription of *mtlA2* than in the absence of inducing cCF10 (Table 3). qRT-PCR results showed that *mtlR* expression in the in the absence of cCF10 induction was four-fold higher than from its natural chromosomal context in OG1RF (pCIE) in both M9YEG and M9YEM media; cCF10 induction led to a further three-to four-fold induction on both media (Table 4). Therefore, increased *mtlA2* expression in mannitol was not due to a difference in *mtlR* levels, but presumably due to mechanisms of carbon catabolite repression (CCR) in glucose.

**Table 3.** *mtlA2* levels in OG1RFΔ*mtlR* complemented strains

| Plasmid | Glucose | Mannitol |
| --- | --- | --- |
| pCIE vector only | 0.032±0.005 | 0.060±0.009 |
| pCIE::*mtlR*+ | 0.225±0.088 | 2.042±0.936 |
| pCIE::*mtlR*+ plus cCF10 <sup>a</sup> | 1.354±0.120 | 2.762±0.427 |
| pCIE::*mtlR* H334A | 0.035±0.006 | ND |
| pCIE::*mtlR* H334D | 0.109±0.007 | 1.306±0.101 |
| pCIE::*mtlR* H392A | 0.029±0.002 | ND |
| pCIE::*mtlR* S412A | 0.613±0.132 | 3.035±0.135 |
| pCIE::*mtlR* H586A | 1.914±0.126 | ND |
| pCIE::*mtlR* H586D | 1.434±0.085 | ND |

<sup>a</sup> cCF10 was added to culture at a concentration of 5 ng/ml 1 hour prior to harvest.

**Table 4.** *mtlR* levels in OG1RFΔ*mtlR* complemented strains

| Plasmid | Glucose | Mannitol |
| --- | --- | --- |
| pCIE vector only | 0.244±0.034 | 0.287±0.057 |
| pCIE::*mtlR*+ | 0.917±0.137 | 1.240±0.344 |
| pCIE::*mtlR*+ plus cCF10 <sup>a</sup> | 3.510±0.464 | 3.817±0.561 |

<sup>a</sup> cCF10 was added to culture at a concentration of 5 ng/ml 1 hour prior to harvest.

The *B. subtilis* MtlR protein is a multi-domain transcriptional activator whose function is regulated by phosphorylation at multiple residues (27). It consists of an N-terminal DNA binding domain, two consecutive PTS regulatory domains (PRD1 and PRD2), an EIIB^Gat^-like domain, and an EIIA^Mtl^-like domain (Fig. 5). The PRD2 domain performs a CCR function by sensing the phosphorylation state of general PTS protein Hpr. In the presence of a preferred carbon source, Hpr dephosphorylates two histidine residues in PRD2, which inhibits MtlR function. In contrast, dephosphorylation of an EIIB^Gat^-like domain cysteine residue and an EIIA^Mtl^-like domain histidine residue by the mannitol-specific EIIA and EIIB components, respectively, activates MtlR in response to the presence and transport of mannitol. In the case of *B. subtilis* MtlR, the phosphorylation state of the cysteine residue is most important. The *E. faecalis* MtlR is 28% identical and 51% similar to *B. subtilis* MtlR and shows homology through all five domains. In particular, all of the relevant histidine residues are conserved in the PRDs and the EIIA^Mtl^-like domain (Fig. 5). However, in *E. faecalis* a potentially phosphorylatable serine residue replaces the *B. subtilis* MtlR EIIB^Gat^-like domain cysteine residue. To determine if these residues are functionally conserved, non-phosphorylatable alanine and phosphomimetic aspartic acid replacements were made in residues analogous to those previously examined in *B. subtilis* MtlR. The mutants were then used to complement the Δ*mtlR* strain from pCIE without cCF10 induction. Results are shown in Table 3. Alanine replacement of the PRD2 histidine residues (H334A and H392A) inactivated MtlR as was observed in *B. subtilis*, suggesting that it performs a similar CCR function. Substitution with aspartic acid at H334 (H334D), however, did not lead to constitutive activity, suggesting either that both histidines in the *E. faecalis* MtlR must be phosphorylated or that the aspartic acid is not sufficiently phosphomimetic in this context. In support of the former interpretation, growth on mannitol led to transcriptional activation of *mtlA2* by the H334D MtlR mutant presumably due to phosphorylation of H392. Replacement of the EIIB^Gat^-like domain serine with alanine (S412A) resulted in partial derepression of *mtlA2* in glucose but did not have as dramatic effect as the cysteine to alanine substitution in *B. subtilis* MtlR. The S412A mutant could be further activated by growth in mannitol indicating the importance of other residues. Replacement of the EIIA^Mtl^-like domain histidine with either alanine or aspartic acid (H586A and H586D, respectively) resulted in full activation of *mtlA2* expression on glucose. Constitutive activation was expected for the alanine mutation, but the aspartic acid substitution in *B. subtilis* MtlR resulted in inactivation of the protein. It is possible that aspartic acid is not sufficiently phosphomimetic in the *E. faecalis* MtlR context.

**Figure 5.**
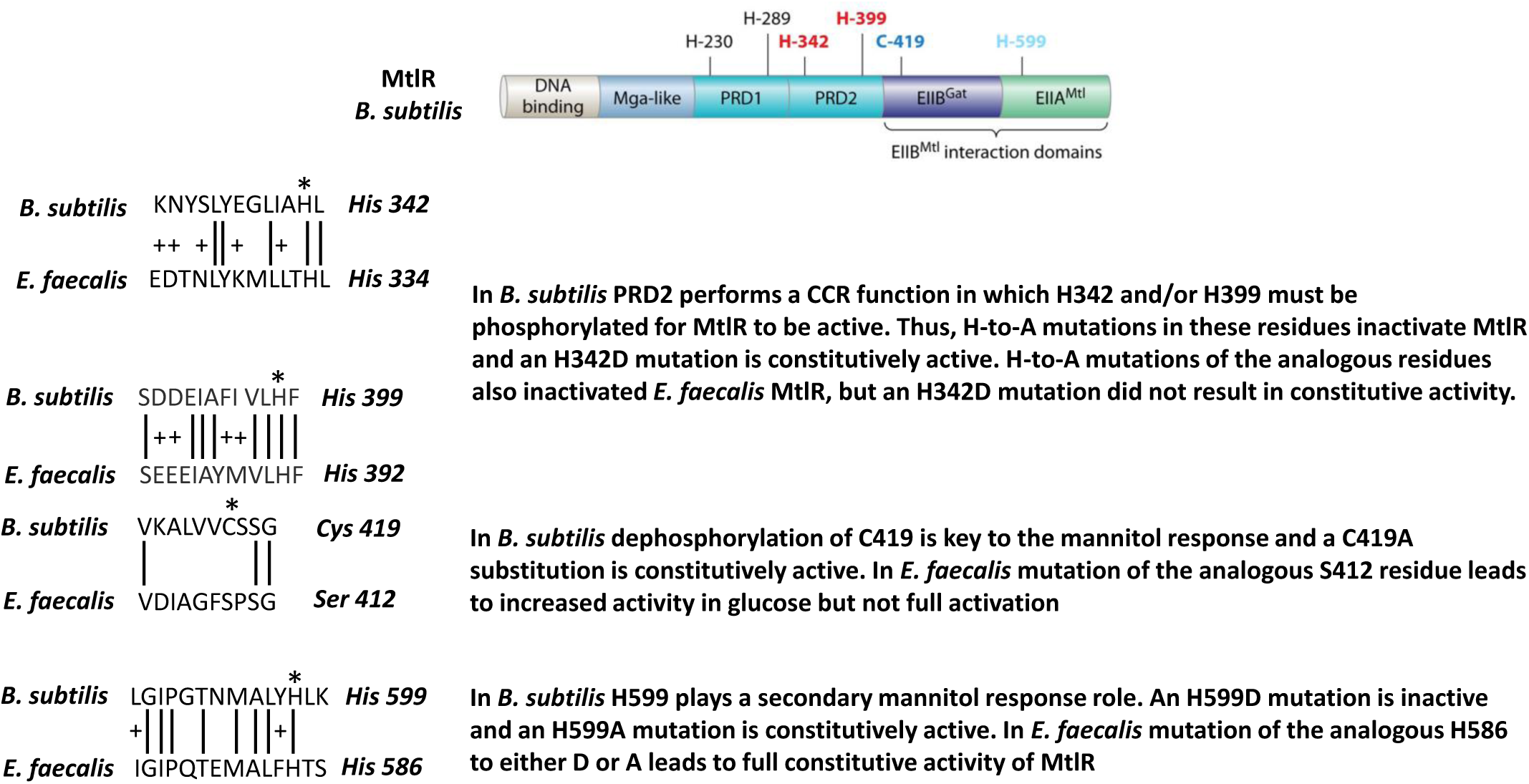
Comparison of *B. subtilis* and *E. faecalis* MtlR. *B*.*subtilis* domain structure with key phosphoylatable residues indicated in color is shown above (reprinted with permission from (26). *E. faecalis* has a similar domain structure with the analogous phosphorylatable residues and surrounding homology shown on the left. The effects on mutations in these residues are compared in the text on the right

As shown in Fig. 1, a putative MtlR binding site was identified upstream of the *mtlA2* promoter based on homology to known sequence in *B. subtilis* (28). Strains with a scrambled sequence in this site phenocopied the Δ*mtlR* mutant. Thus, the MtlR binding site mutation was unable to grow on M9YEM (Fig. 2B) and showed reduced levels of the *mtlA2* transcript compared to WT on M9YEG (Table 2). Provision of excess MtlR ectopically did not complement the binding site mutation (Fig. 2B).

Another potential candidate for a CCR role in regulating the *mtlA2* operon is catabolite control protein A (CcpA). CcpA functions to repress genes involved in the metabolism of non-preferred carbon sources in the presence of glucose. Indeed, a *ccpA* deleted strain (kindly provided by the Garson lab) showed elevated expression of *mtlA2* in M9YEG and an even greater level of derepression in M9YE with 90% glucose and 10% mannitol (Table 5). Repression was restored in a complemented strain. A putative CcpA binding site (CRE) was identified, based on sequence homology to the CRE site of the *E. faecalis eutS* gene (29), between the -10 and -35 promoter elements of the *mtlA2* operon (Fig. 1). Unfortunately, mutants constructed in this element eliminated *mtlA2* expression, so a more detailed examination of the sequence will be required to confirm its function.

**Table 5.** *mtlA2* levels in *ΔccpA* and complemented strains

| Strain | Glucose | 90%Glu+10%Mtl |
| --- | --- | --- |
| WT | 0.228±0.046 | 0.959±0.082 |
| $\Delta ccpA$ | 0.314±0.023 | 2.092±0.075 |
| $\Delta ccpA::ccpA+$ | 0.187±0.006 | 1.024±0.092 |

With the function of the *mtlA2* operon and the *mtlR* gene established, the question of the purpose of the *mtlAF* paralogous PTS system remained. In both *B. subtilis* and *L. lactis*, MtlR is negatively regulated by phosphorylated MtlF and *mtlF* mutations lead to derepression of the mannitol transporter in glucose. To investigate if the *E. faecalis mtlAF* PTS might perform a similar regulatory function, in-frame deletions of the two components were tested for impact on *mtlA2* expression. Consistent with a negative regulatory role, deletions resulted in a two-three fold increase in *mtlA2* expression in M9YEG compared to WT (Table 2). The deletions had no effect on *mtlA2* expression in M9YEM.

### Between the operons, what is *par*_EF0409_ doing?

In most published genomes of the Firmicutes the genes for mannitol transport, regulation, and metabolism are encoded in a single operon in the order *mtlA-mtlR-mtlF-mtlD*. Variations include the independent transcription of *mtlD* in *Lactococcus lactis* and the separation of *mtlR* from the rest of the operon in *Bacillus subtilis* (28, 30). Using the phylogeny of Zhong et al (31), we searched for *mtlA, mtlA2*, and *mtlD* sequences in enterococcal species by NCBI blastn. Twenty-five of the 34 species in the dendogram lacked homologs to all three genes and therefore are unlikely to be able to utilize mannitol as a carbon source. Of the nine remaining species, *Enterococcus gilvus, Enterococcus avium*, and *Enterococcus raffinosus* encode an *mtlA-mtlR-mtlF-mtlD* operon. The remaining six species (*E. faecalis, E. faecium, Enterococcus mundtii, Enterococcus thailandicus, Enterococcus casseliflavus* and *Enterococcus gallinarum*) encode both an *mtlA-mtlR-mtlF* and an *mtlA2-mtlF2-mtlD* operon but with significant variability in spacing and intervening genes. The *par*_EF0409_ TA-1 locus is unique to *E. faecalis* and none of the other species encode homologs of *par* Fst toxins or evidence of a TA locus. *E. mundtii* is the most divergent with the two operons separated by 14.5 Kbp and multiple genes. In *E. faecium* the operons are about 2.3 Kbp apart and the ATP binding and permease components of an ABC transporter are encoded in the gap and transcribed in the opposite direction to the mannitol operons. The closely related species *E. casseliflavus* and *E. gallinarum* have similar inter-operon gaps of 313 nt, significantly shorter than the *E. faecalis* 611 nt gap. The gap includes two inverted repeats that show characteristics of intrinsic terminators, but no direct repeats or open reading frames. *E. thailandicus* has an inter-operon gap of 544 nt and a complex combination of direct and inverted repeats, but with different spacing than *par*-related TA systems and no apparent open reading frame.

Whether the *E. faecalis par*_EF0409_ TA locus plays a unique adaptive role or is merely an evolutionary accident, e.g., the remnant of a long lost MGE, is not clear. To examine whether there might be a link between *par*_EF0409_ function and mannitol metabolism, we compared the toxicity of Fst_EF0409_in M9YEG and M9YEM. As show in Fig. 6, cells grown in M9YEM were approximately 10-fold more sensitive to toxin expression than cells grown in M9YEG as shown by similar growth inhibition at 5 ng/ml inducing cCF10 and 0.5 ng/ml cCF10 in M9YEG and M9YEM, respectively. A similar increase in sensitivity on mannitol was observed when cellular viability was assessed (data not shown).

**Figure 6.**
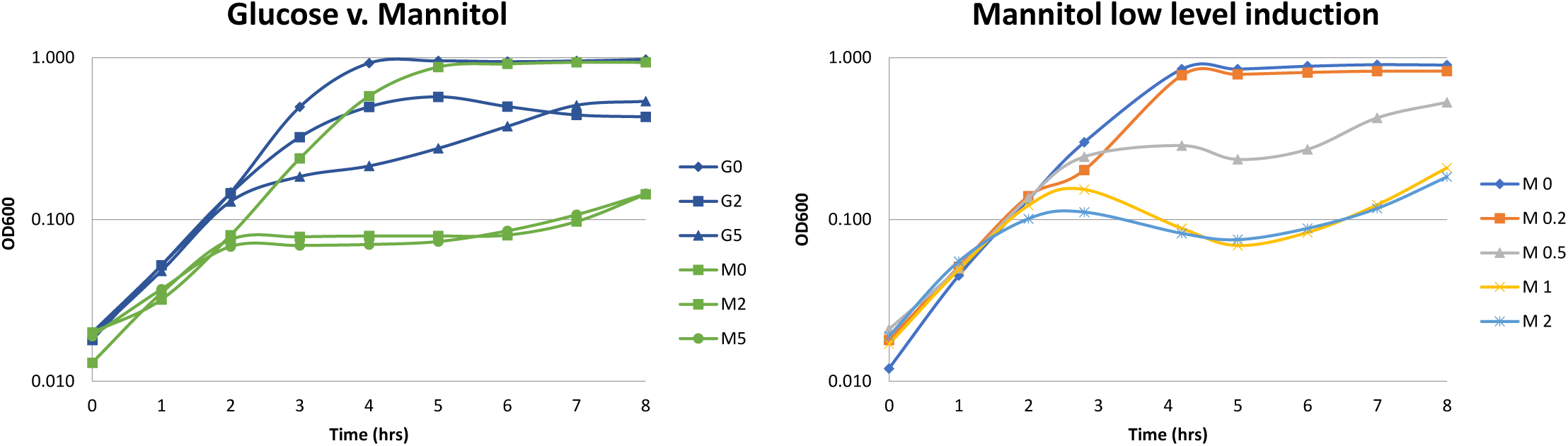
Toxicity of Fst_EF0409_ TA-1 toxin in cells grown in M9YEG vs. M9YEM. Fst_EF0409_ was expressed from the cCF10 responsive P_Q_ promoter of pCIE. G=glucose. M=mannitol. Numbers are the concentration of cCF10 added in ng/ml. The graph on the left is a comparison of growth in glucose and mannitol at the same inducing cCF10 levels. The graph on the right shows the effects of lower levels of inducing cCF10 on cells grown in mannitol.

One potential function for *par*_EF0409_ might be to suppress recombination between homologous regions of the *mtlA* and *mtlA2* genes. If such recombination occurred, *par*_EF0409_would be on the excised DNA. Its loss during outgrowth of recombined strains would trigger TA-1 PSK function. Comparison of the *mtlA* and *mtlA2* sequence revealed several regions of significant homology at which recombination could occur. To detect recombinants, primers were synthesized complementary to regions of non-homology flanking the homologous regions. PCR was then performed and products examined for the predicted size of recombinants. In addition, primers were synthesized to amplify across the junction of the predicted excised circle. Products of the appropriate size were detected with both primer sets using either WT OG1RF or an isogenic strain deleted for *par*_EF0409_ (data not shown). PCR products were cloned in pGEMTeasy and sequenced to identify the junction sites. The majority of recombination events occurred within a 54 nt region with 85% identity between nucleotides 306486-306539 in *mtlA* and 311329-311382 in *mtlA2* encoding the conserved P-loop motif in the IIB domain. In particular, a 5’-ATGGGGGC-3’ motif within this region was a hotspot for recombination. The same bias was observed in WT and mutant strains and in predicted chromosomal and excised circle recombinants.

While these results are suggestive of a link between TA function and the function of the surrounding genes, interpretation requires several caveats. As with any over-expression study, the relevance to genes in their normal context must be considered. In all conditions examined so far, the antitoxin RNA is produced in large molar excess over the toxin message (9). Results from this study suggest this is also true in mannitol. RNA-seq experiments showed that the toxin-encoding message, RNA I_EF0409_ (designated OG1RF_RS01630 in supplemental data), is barely detectable in both M9YEG and M9YEM. RNA I_EF0409_ was detectable by qRT-PCR and Northern blot, but levels were low and variable and no significant differences between growth media were observed (data not shown). As shown in Fig. 7, RNA II_EF0409_ antitoxin levels were reduced in M9YEG in stationary phase relative to M9YEG in Northern blots. However, log phase cultures showed no difference in antitoxin levels. Therefore, although over-expression data suggests that mannitol-grown cells may be more sensitive to the toxin than glucose-grown cells, conditions under which expression of RNA I_EF0409_ from its natural position would be high enough to escape RNA II_EF0409_-mediated repression have yet to be identified.

**Figure 7.**
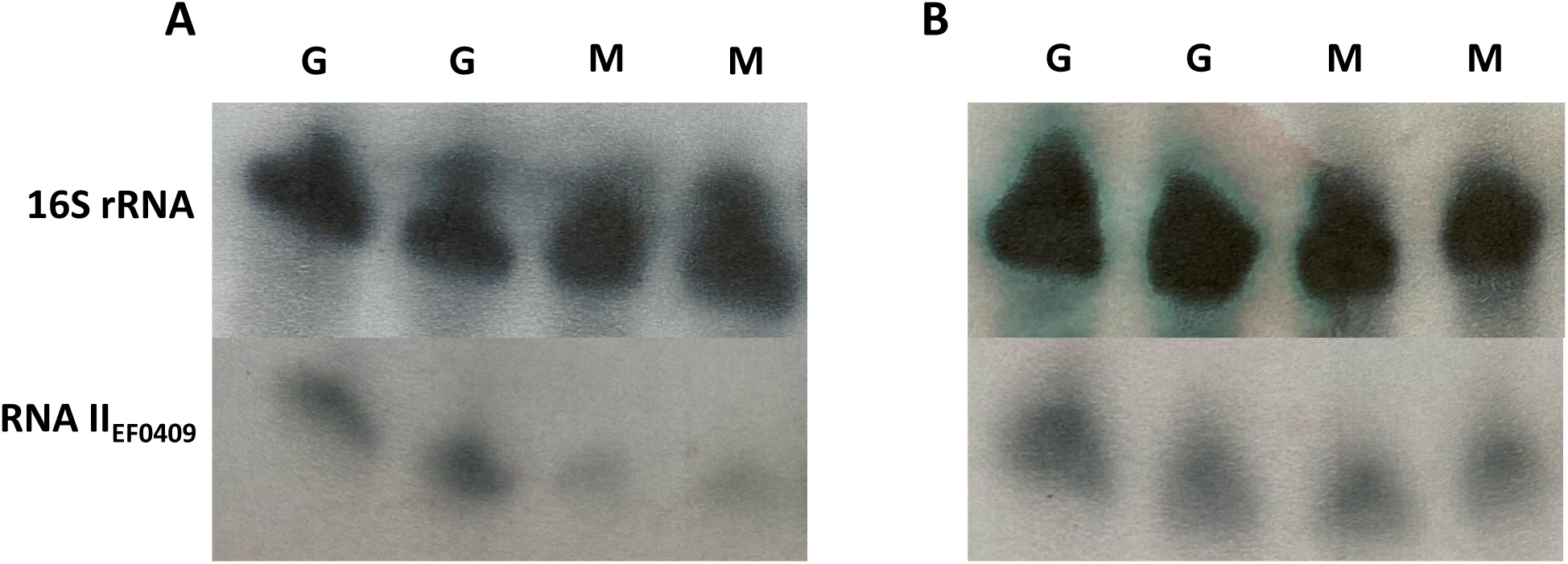
RNA II_EF0409_ levels in glucose- and mannitol-grown cells. 16S rRNA was used as a loading control. G = M9YEG-grown and M = M9YEM grown. Duplicate samples were performed with RNA from independently grown cultures. **A**. Stationary phase. **B**. Log phase.

## Discussion

In this report, we define the genetic organization and functional roles of paralogous mannitol-family PTS systems in *E. faecalis* strain OG1RF. The relevant genes are organized into two operons flanking the *par*_EF0409_ TA-1 system. The downstream *mtlA2* operon is highly induced by mannitol and encodes the essential transport apparatus, *mtlA2F2*, and metabolic enzyme, *mtlD*, for mannitol utilization. This operon was partially described previously (8). The upstream *mtlA* operon is constitutively transcribed and encodes genes essential for proper regulation of the *mtlA2* operon. This includes the *mtlR* gene, which is homologous to mannitol-responsive positive transcriptional regulators in other organisms and was shown to be essential for mannitol-dependent induction of the *mtlA2* operon. Mutations in *mtlR* also showed reduced basal transcription of *mtlA2* and a slightly reduced growth rate in glucose. The PTS system encoded with *mtlR* is not essential for growth in mannitol, but is required for full repression of the *mtlA2* operon in its absence. While the *mtlAF* genes and show greatest homology to mannitol transporters, we cannot rule out at this time the possibility that *mtlAF* transports some other sugar.

The *E. faecalis mtlR* gene encodes a protein closely related to the well-studied *B. subtilis* mannitol PTS regulator *mtlR*. The effects of mutations in phosphorylatable residues of *E. faecalis* MtlR were generally consistent with those previously observed in *B. subtilis* MtlR (28) with some subtle differences. Thus, alanine substitutions of the two histidine residues in the *E. faecalis* MtlR PRD2 domain resulted in loss of *mtlA2* operon induction in mannitol, supporting a CCR function of this domain. However, unlike in *B. subtilis*, substitution of the phosphomimetic aspartic acid residue at H334 (analogous to H342) did not result in constitutive activation suggesting either that the aspartic acid is not sufficiently phosphomimetic in *E. faecalis* MtlR or that both PRD2 histidines must be phosphorylated. Alanine substitutions in phosphorylatable residues in the *E. faecalis* MtlR EIIB^Gat^-like and EIIA^Mtl^-like domain resulted in constitutive expression of the *mtlA2* operon, suggesting that these two residues are dephosphorylated by the PTS system in the presence of mannitol, as in *B. subtilis* MtlR, to induce the *mtlA2* operon. In *B. subtilis*, the C419 residue in the EIIB^Gat^-like domain plays the predominant role. In *E. faecalis*, this residue is substituted by S412 in and the H586 residue in the EIIA^Mtl^-like domain appears to play the predominant role. One caveat to the interpretation of these results is that complementation from the pCIE vector resulted in a 3-to 4-fold higher expression of *mtlR* mRNA than from the gene in the chromosome (Table 4). It is unclear what effect this difference in transcript dosage might have. A putative MtlR binding site showing substantial sequence similarity to that for *B. subtilis* MtlR was found upstream of the *mtlA2* promoter and mutation of this site prevented mannitol-dependent transcriptional activation of the *mtlA2* gene supporting its role as the site of MtlR action.

CcpA and the MtlAF PTS transporter appear to play secondary roles in repressing the *mtlA2* operon in the presence of glucose and in the absence of mannitol, respectively. CcpA also plays a secondary role to phosphorylation of MtlR PRD2 in repression of mannitol metabolic genes during CCR in other Gram-positive species (32). A putative CRE site for CcpA binding was identified between the -10 and -35 boxes of the *mtlA2* promoter based on homology to a CRE site in the *E. faecalis eut* operon (29) but we were unable to confirm that this site was essential for CcpA function. MtlAF presumably functions through MtlR phosphorylation but the rationale for having a second transporter system is unclear. In other systems, a single transporter appears to be sufficient for this mode of regulation. It is possible that CcpA and/or MtlAF may play greater roles under different culture conditions, e.g., different combinations of carbon sources or synthetic vs. semi-synthetic media.

The distribution of mannitol utilization genes on the enterococcal family tree is sparse and spotty and the operon organization is highly variable. This suggests that the relevant genes were acquired by horizontal gene transfer independently on multiple occasions. Most, but not all, enterococcal species that harbor genes for mannitol transport and metabolism have two presumptive operons similar to the *E. faecalis mtlA* and *mtlA2* operons, but the inter-operon distance varies from 14.5 Kbp and 313 bp. Only *E. faecalis* encodes a discernable TA system within this intervening region. The origin and function of the *par*_EF0409_ TA-1 system remains unclear, but several pieces of circumstantial evidence suggest that its function may be coordinated with that of surrounding genes. First, as previously reported (33), mutation of an efflux pump located two Kbp downstream of *mtlD* results in hypersensitivity to Fst_EF0409_, the *par*_EF0409_ toxin. Located between the *mtlA2* operon and the genes for the efflux system are genes annotated as NAD binding proteins that could conceivably be involved in recycling the NAD+ cofactor required for MtlD function. Second, transcription initiated from the *mtlA* promoter appears to read through into the RNA II_EF0409_, the *par*_EF0409_ antitoxin, potentially impacting its expression and or interaction with RNA I_EF0409_, the toxin-encoding mRNA. Third, mannitol-grown cells are hypersensitive to Fst_EF0409_ expression. Fourth, homologous sequences in *mtlA* and *mtlA2* are prone to recombination, potentially excising the intervening DNA. Since the excised DNA would include the *par*_EF0409_ locus, the TA function might prevent its loss. Why this would be advantageous to *E. faecalis* but not to those species with the two operons separated by similar inter-operon distances is not clear. In addition, hypersensitivity in the efflux system mutant and in mannitol-grown cells was only detected using ectopic overexpression of the toxin. In the natural context, under all growth conditions examined so far, antitoxin is produced in substantial molar excess to toxin mRNA so it is unclear when the cell would be exposed to sufficient toxin to observe an effect. The data presented here provides essential context for further efforts to determine the function of *par*_EF0409_ as well as to examine the evolution of carbohydrate metabolism in the enterococci.

## Materials and Methods

### Bacterial Strains, Media, and Growth Conditions

*E. faecalis* strain OG1RF was used for all experiments in this study (34, 35). An OG1RF strain deleted for *par*_EF0409,_ OG1RFΔ*par*_EF0409_ (9), was also used in recombination studies. Transposon mutant strains were obtained from the ordered library constructed in OG1RF by the G. Dunny laboratory at the University of Minnesota (22). The location of transposon inserts based on the NCBI OG1RF genome sequence are as follows: *mtlA* (305429), *mtlR* (308259), *mtlF* (309542), *mtlA2* (310258), *mtlF2* (312088 and 312328). For growth curves and RNA preparation *E. faecalis* cultures were grown in M9YE base (36) supplemented with either 20 mM glucose or mannitol. *E. faecalis* liquid cultures were grown at 37 °C with shaking at 25 rpm. Agar plates were prepared by addition of 17g agar (Research Products International, Mt. Prospect, IL, USA) per one liter of medium. Plates were incubated overnight at 37 °C. Chloramphenicol (Cm) (Sigma-Aldrich, St. Louis, MO, USA) was added for plasmid selection where appropriate. Overnight cultures were grown with 25 µg Cm per ml and diluted to one or two percent in fresh medium with 10 µg Cm per mL. Expression from pCIE was established by the addition of peptide pheromone cCF10 (H-LVTLVFV-OH) (Mimotopes Clayton, Australia) at concentrations indicated for individual experiments as previously described (9). An equal volume of solvent (dimethylformamide) was added for uninduced controls. Plasmid DNA was introduced into *E. faecalis* by electroporation as previously described (37). The *E. coli* strain DH5α (New England Biolabs, Ipswich, MA, USA) was used for sub-cloning procedures and plasmid purification. *E. coli* cultures were grown in Luria-Bertani (38) medium with antibiotics at the following concentrations where appropriate: Ampicillin (amp), 100 µg/mL; Cm, 25 µg/mL. *E. coli* liquid cultures were grown at 37 °C with shaking at 250 rpm.

### Genetic Manipulations

In-frame, markerless deletions of *mltA, mtlR, mtlF* and *mltA2* were constructed in OG1RF by allelic exchange using vector pJH086 (39) as previously described (33, 40). The mutant alleles were synthesized by Blue Heron Biotech, LLC, (Bothell, WA, USA) and contained the first 5 and last 5 codons of the corresponding gene and approximately 900 base pairs of flanking DNA. The construct was synthesized with *PaeI* and *Smal* restriction sites at each end, and these enzymes were used to subclone the fragment from the commercially provided pUCminusMCS vector to pJH086. After a post-ligation cut with *BamH*I, plasmids were purified and introduced into competent DH5α cells with selection for Cm and growth at 30°C. Restriction enzymes and DNA ligase were obtain from New England Biolabs, Ipswitch, MA, USA. Plasmids were purified by QIAprep Spin Miniprep kit (Qiagen, Valencia, CA, USA), checked for appropriate restriction pattern, and introduced into *E. faecalis* cells for allelic exchange. Recombinants were screened by colony PCR (41), using primers flanking the desired deletion (Table S1). PCR products showing the appropriate size for the deletion were then sequenced to ensure that no spurious mutations were obtained. All primers were produced by Integrated DNA Technolgies (IDT, Coralville, IA, USA) All DNA sequencing was performed by Eurofins Genomics LLC (Louisville, KY, USA).

Mutations in the putative MtlR and CcpA (CRE) binding sites were constructed in a similar manner to the deletions with the exception that synthesized sequences contained scrambled binding site sequences flanked by approximately 900 base pairs for recombination. The MtlR binding site sequence was changed from TTTGGCACAATGGTTTGTGCCAA to ACATGGCTCGATTAGATGTATGTC and the CRE site was changed from GGAAAGC to GACAGGA. Allelic exchange was carried out as above. In the case of the MtlR binding site mutants, roughly half of the isolated recombinants showed a growth defect on M9YEM plates. These recombinants were confirmed to contain the desired mutation by PCR using flanking primers SMBS-F and SMBS-R (Table S1) and sequencing. The recombinants containing the scrambled CRE site were identified by PCR using the SCRE-detect primer that binds to the scrambled sequence and SCRE-R primer flanking the region. Presence of the scrambled CRE sequence was confirmed by sequencing the product amplified using SCRE-F and SCRE-R primers (Table S1).

For complementation studies, a promoterless version of the *mtlR* gene was amplified by PCR from OG1RF genomic DNA purified as previously described (41) using primers indicated in Table S1. Amplified product and pCIE were digested with *BamHI* and *SphI*. After a post-ligation cut with *Sal*I plasmids were purified and introduced into competent DH5α cells with selection for Cm and growth at 37°C. This construct, designated pCIE::*mtlR*+, was then used as template for introduction of the relevant MtlR mutations by GenScript (Piscataway, NJ, USA). These constructs were then introduced into OG1RFΔ*mtlR* by electroporation.

### RNA purification and analysis

RNA samples were prepared for all purposes as previously described (9). 5’Rapid Amplification of c-DNA Ends (5’-RACE) was performed using the Invitrogen 5′ RACE System for Rapid Amplification of cDNA Ends, Version 2.0 (Carlsbad, California) according to manufacturer’s instructions with modifications as described in (42). Quantitative reverse transcription-PCR (qRT-PCR) was performed as previously described (9). All data represents the average of at least three biological replicates. Northern blots were performed as previously described (43) with oligonucleotide probes specific for RNA I_EF0409_, RNA II_EF0409_and *E. faecalis* 16S rRNA as a loading control as in (9). RT-PCR for identification of polycistronic RNAs was performed as previously described (9) with primer pairs shown in Table S1. The PCR temperature cycling conditions were as follows: initial denaturation for 1min at 94°C, followed by 30 standard cycles: denaturation at 94°C for 30 sec, primer annealing for 30 sec at 55°C, and primer extension at 72°C for 1 min. The last cycle was followed by 10 min incubation at the primer extension temperature of 72°C. Size of the products were checked by running them on an agarose gel. Products were sequenced by Eurofins Genomics LLC (Louisville, KY, USA) using the gene specific primers (Table) to ensure it is not a spurious product from some other region.

RNA-seq was performed by the University of Nebraska DNA Sequencing Core at the University of Nebraska Medical Center in Omaha, NE. Libraries were constructed from 300 ng of total RNA from sample provided by our laboratory using the TruSeq® Stranded Total RNA Library Prep (Illumina, Inc. San Diego, CA) following the recommended protocol. In the step in which the total RNA is incubated with Ribozero to remove ribosomal transcripts, 5 L of bacterial ribozero solution was used since these cultures do not have contaminating eukaryotic RNA. Resultant libraries from the individual samples were multiplexed and subjected to 75 bp paired read sequencing to generate approximately 60 million pairs of reads per sample using a HighOutput 150 cycle flowcell on an Illumina NextSeq500 sequencer in the UNMC Genomics Core facility. The original fastq format reads were trimmed by fqtrim tool (https://ccb.jhu.edu/software/fqtrim) to remove adapters, terminal unknown bases (Ns), and low quality 3’ regions (Phred score < 30). The trimmed fastq files were processed by FastQC (44). The reference genome for *Enterococcus faecalis* OG1RF was downloaded from Ensembl. The trimmed fastq files were mapped to *Enterococcus faecalis* by CLC Genomics Workbench 12 for RNAseq analyses. For statistical analysis, each gene’s read counts are modeled by a separate Generalized Linear Model (GLM), assuming that the read counts follow a negative binomial distribution and were normalized based on transcripts per million (TPM). The Wald test was used for statistical analysis of the two-group comparisons. The false discovery rate (FDR) and Bonferroni adjusted p values were also provided to adjust for multiple-testing problem. Fold changes are calculated from the GLM, which corrects for differences in library size between the samples and the effects of confounding factors. Raw and processed files are available at the GEO repository: accession number GSE195652.

### Detection of *mtlA*-*mtlA2* genomic recombinants

Genomic DNA was purified as previously described (41). Primers were designed to read across regions of homology between *mtlA* and *mtlA2* at which recombination was predicted to occur (Table S1). Depending upon the combination of primers used in PCR, products would be produced from genomic recombinants (5’-*mtlA* + *mtlA2*-3’), the predicted excised circle (5’-*mtlA2* + *mtlA*-3’), or the unrecombined genes (5’*-mtlA* + *mtlA*-3’ and 5’-*mtlA2* + *mtlA2*-3’). PCR was performed using Kapa Biosystems KAPA2G Fast PCR mix according to the manufacturer’s instructions (Wilmington, MA, USA). PCR products were cloned into pGEM-T Easy and transformed into DH5α competent cells (Promega). Plasmid DNA was purified using the Qiagen midiprep kit (Qiagne, Valencia, CA, USA) and DNA was sequenced at Eurofins Genomics.

## Acknowledgements

This research was funded by PHS grant AI140037. RNA-seq was performed at the University of Nebraska DNA Sequencing Core. The Core receives partial support from the National Institute for General Medical Science (NIGMS) INBRE - P20GM103427-19 grant as well as The Fred & Pamela Buffett Cancer Center Support Grant - P30 CA036727. This publication’s contents are the sole responsibility of the authors and do not necessarily represent the official views of the NIH or NIGMS.

